# Counting apples - How does the prefrontal region support early numerical understanding in preschool children?

**DOI:** 10.64898/2026.01.30.702872

**Authors:** Elizaveta Ivanova, Emily K Farran, Mojtaba Soltanlou

## Abstract

Because early maths skills strongly predict later outcomes, it is crucial to understand the mechanisms that shape early learning in children. The recent years have seen an increase in studying the neural correlates that support the acquisition of maths skills. However, existing work in early childhood has primarily focused on core number-processing regions in the parietal regions, with comparatively little attention to the supportive role of prefrontal regions. In this study, we examined the engagement of the prefrontal regions when matching numbers and objects. Children (N=60, 25 girls, aged 2.74-5.18 years) matched auditory small (1-3) and large (5-7) numbers, as well as objects (fruits) to corresponding visual pictures while their frontoparietal brain responses were recorded using functional near-infrared spectroscopy (fNIRS). Importantly, matching large numbers was substantially more difficult than matching small numbers or objects. The analysis revealed that children had increased activation in the right middle frontal gyrus when matching large numbers, compared to small numbers. However, there was no difference in the prefrontal region between matching small numbers and objects. The connectivity analysis further revealed increased frontoparietal connectivity when matching small numbers, but not large numbers or objects. Our findings suggest that prefrontal involvement during early numerical knowledge acquisition relies primarily on domain-general mechanisms, with number-specific responses likely to emerge later in development.

## Introduction

The acquisition of symbolic numerical knowledge marks a major milestone in early cognitive development, laying the foundation for later mathematical achievement (Hyde, 2021). In contrast to non-symbolic numerical knowledge, which is evident within the first months of life (Izard et al., 2008; Hyde et al., 2010; Edwards et al., 2016), understanding of numerical symbols typically emerges between the ages of three and five years (Sarnecka et al., 2015). Notably, earlier acquisition of symbolic numerical knowledge predicts later mathematical development (Geary et al., 2018), underscoring the importance of identifying the neural mechanisms that support this developmental cascade.

Neuroimaging evidence suggests that both parietal and prefrontal regions contribute to numerical cognition in early childhood (Hyde, 2021). The parietal cortex, particularly the intraparietal sulcus (IPS), has been most consistently associated with numerical processing. The right IPS supports magnitude representations from infancy, allowing discrimination of numerical quantities based on ratio-dependent sensitivity: the smaller the ratio (e.g., 6:7 compared to 1:2), the stronger the activation due to increased processing demands (Ansari & Dhital, 2006; Holloway & Ansari, 2010; Piazza et al., 2004; Piazza et al., 2007; Halberda & Feigenson, 2008; Dehaene, 2011). Importantly, sensitivity to numerical processing in the right IPS appears broadly comparable across preschoolers, older children, and adults (Cantlon et al., 2006; Kersey & Cantlon, 2017). In contrast, the left IPS becomes increasingly engaged as symbolic numerical knowledge develops (Cantlon et al., 2006; Seghier et al., 2013), with left-lateralised number-specific responses emerging between approximately three and five years of age (Pinhas et al., 2014; Park et al., 2014; Zhang et al., 2019). These findings suggest that numerical processing in early childhood involves both hemispheres of the parietal cortex, with the right IPS providing support early on and the left IPS becoming increasingly involved around the time preschoolers start gaining symbolic numerical knowledge.

Surprisingly, much less is known about the contribution of the prefrontal regions in the development of symbolic number processing in early childhood. There are two views on the role of prefrontal regions in numerical cognition. According to a well-established view, prefrontal engagement in early number processing in preschoolers is mainly domain-general, and as such can only be modulated by cognitive load via task difficulty or competency (Ayres, 2006). To this end, Cantlon et al. (2009) found that when comparing numbers, children aged 6-7 years old recruited the inferior frontal cortex to a greater extent than adults, as the task was more difficult for them. Similarly, Battista et al. (2018) reported a longitudinal decrease in connectivity between parietal and prefrontal regions in children aged 7-14 years when judging the correctness of simple arithmetic problems (e.g., 2+4=6), suggesting lesser reliance on prefrontal support. Simon et al. (2015) observed in 5-7-year-old children an increase in the anterior cingulate cortex when presented with a conflict version of the number conservation task (e.g., five objects in two rows, but in one row the space between objects is much larger, making the set of objects look visually longer), but not when presented with congruent trials that did not increase cognitive load. Taken together, there is strong evidence that prefrontal engagement in preschoolers and older children is largely modulated by cognitive load across a wide range of tasks that require symbolic numerical knowledge.

The second view posits that the prefrontal cortex, as part of the frontoparietal network supporting mathematical thinking, may also exhibit number-specific responses early on. Under this view, prefrontal regions show a preference for numerical over non-numerical processing when task difficulty is matched. Studies in older children and adults, but not preschoolers, have found supporting evidence for number specificity. Using a domain-matched comparison task involving faces, words, shapes, and numbers, designed to be similarly easy across conditions, Emerson and Cantlon (2012) reported number-specific responses in the inferior frontal gyrus and insula in both children aged 4-11 years and adults. Using the same paradigm, Emerson and Cantlon (2014) further identified number-specific prefrontal activations in bilateral anterior cingulate cortex, inferior frontal gyrus, and contiguous insular cortex in children aged 4-9 years as well as in adults. Approximately one-third of these activations overlapped with activity modulated by reaction time across tasks, likely reflecting the recruitment of domain-general resources under increased cognitive load. As such, Emerson and Cantlon’s findings suggest that symbolic number processing engages both number-specific and domain-general prefrontal mechanisms. Koyama et al. (2017) found that in adults, despite the functional overlap in the middle frontal gyri (MFG), left MFG is more engaged during literacy-related tasks, and right MFG is associated with numerosity-related tasks. Finally, in an audio-visual matching task, Bugden et al. (2021) observed activation in the anterior cingulate cortex only in adults, but not in 4-10-year-old children. As both age groups’ performance was at the ceiling effect, the resulting prefrontal activation in adults could be attributed to a number-specific response, while the lack of such activation in children could suggest that the number-specific prefrontal response was yet to be developed.

Taken together, existing evidence for number-specific prefrontal engagement is largely limited to older children and adults, leaving open the question of when such specificity emerges developmentally. If prefrontal involvement in early symbolic number processing is purely domain-general, this would underscore the crucial contribution of broad cognitive skills to the acquisition of numerical symbols. Conversely, if number-specific prefrontal engagement is present from early on, it would imply a specialised pathway for symbolic numerical learning that is less reliant on domain-general resources than previously assumed.

## The present study

Our study aimed to shed light on the role of the prefrontal region in numerical development in preschool children. In particular, it remains unclear whether prefrontal engagement at this developmental stage is only domain-general. If so, it would only stem from the contrast of difficult vs. simple. Alternatively, this engagement may also be number-specific, e.g., differentiated for the number vs. non-number dichotomy. To explore these questions, we used a version of the audio-visual matching task (Bugden et al., 2021; Pinhas et al., 2014), in which children were asked to identify whether the auditorily presented number words matched the number of visually presented objects. Brain responses were recorded using functional near-infrared spectroscopy (fNIRS), a child-friendly neuroimaging method that allows ecologically valid testing (Barreto & Soltanlou, 2022; Soltanlou et al., 2018).

To capture domain-general engagement, we introduced two levels of difficulty. In the easy numerical task, children matched audio-visual pairs containing small numbers ranging from ‘one’ to ‘three’. In the difficult numerical task, the audio-visual pairs consisted of larger numbers ranging from ‘five’ to ‘seven’. To test for number-specific engagement, we included a non-numerical task, similarly easy to do as the easy numerical task, in which children were required to semantically match audio-visual pairs of familiar objects (fruits) without counting. To keep children’s attention across all three levels and avoid automatic responses, stimuli were presented in two formats: matched pairs, where the auditory and visual information corresponded, and mismatched pairs, where they did not align.

We formulated the following main hypotheses: if prefrontal engagement in preschoolers is primarily driven by domain-general support, the activation should increase only when matching large numbers compared to matching small numbers, given the higher cognitive demands associated with larger numerosities. If, however, prefrontal involvement also includes number-specific contributions, activation should also differ between matching small numbers and matching objects.

In addition, we investigated the responses in the bilateral parietal regions, as they play a key role in number processing within the frontoparietal network of mathematical thinking. So as to not shift our focus from the main hypotheses about domain-general and number-specific prefrontal engagement, we formulated broader, less specific expectations for parietal activation: for both difficult vs. simple number matching and simple vs. non-number matching, we expected bilateral parietal regions to elicit increased activation, due to increased processing demands in the former and number specialisation in the latter.

## Materials and Methods

### Participants

Children aged 2.74-5.18 years old (N=60; M=3.94, SD=0.49, 25 girls) participated in this study. All children had no history or diagnosis of chronic neurological or mental illness, visual or auditory impairment, or serious head injury. Due to the possibility of testing children in English and Russian, we invited the children who speak English or Russian as their primary language. As English and Russian languages both require singular-plural marking, and native speakers of both languages develop early numerical skills within the same timeline (Sarnecka et al., 2007), the language will not be a factor in the analyses. Eight children completed the study in the Russian language. Caregivers gave written informed consent and received an expense allowance for their participation. Similarly, children gave verbal consent and received a toy for their participation. All procedures were in line with the latest ethics guidance of the Declaration of Helsinki and were approved by the Ethics Committee of the University of Surrey.

For one child, the recording was restarted during the first block due to technical issues; the initial recording was discarded, and only the second attempt was included in the analyses. For four children, the recording was briefly paused mid-block to allow the child to take a short break. In addition, due to a technical error, the final three trials of the non-numerical task were not recorded for one child. All children were retained in the final analyses. The data of the remaining 59 children was complete.

### Behavioural measures

In addition to the collected demographic information, children completed a series of paper-and-pencil behavioural tasks to characterise their development across numerical and domain-general cognitive domains.

To assess symbolic numerical knowledge, we used the titrated version of the Give-a-Number task (Wynn, 1990; Krajcsi, 2021). In this task, children were given 30 small balls (2 cm in diameter) and were asked to provide a specific number of them. The task was divided into three sets with a pseudorandomised numerical sequence in each: i) 1, 3, 2, ii) 6, 4, 5, iii) 9, 7, 8. The task was presented in three runs, and in each run, a child was always presented first with the set that contained numbers 1, 3 and 2. The progression to the next set was dependent on a child providing correct responses for at least two numbers out of three in the current set. For instance, if a child correctly responded to numbers 1 and 2, but not 3, they would still move on to the next set, within the same run. This was done to account for the potential shyness/distractedness. After three runs of the task, a number was considered known if the correct response for that number was given at least in two out of three runs, and all smaller numbers were scored as known numbers. Consequently, the highest number known was considered the child’s score and was used as the measure of symbolic numerical knowledge.

The Preschool Early Numeracy Screener (PENS, Purpura et al., 2015) was used to measure overall competency in early numeracy. The items covered areas such as verbal counting, numerical relations, and arithmetic operations. The raw score was calculated as the total number of correct responses, stopping after three consecutive incorrect answers. The total raw score was used as the measure of overall early numeracy.

Working memory was assessed using the forward letter span task (Soltanlou et al., 2015). In this task, the child was asked to repeat sequences of consonants. They were first presented with two sequences, each containing two letters. If the child correctly repeated at least two out of the three sequences, they were then presented with the next sequence, which had one additional letter. The maximum number of letters presented was four. The highest number of letters successfully repeated in at least two sequences was considered the child’s final score. This score was used as the measure of verbal working memory.

To assess non-verbal IQ, we administered the standardised, norm-referenced Matrices subtest from the British Ability Scales III (BAS III, Elliot & Smith, 2011). This subtest evaluated non-verbal IQ across various abilities, including visual-perceptual organisation, nonverbal reasoning, and cognitive flexibility. There were 18 items in total, and the number of correctly solved items was calculated as the score of each child. Therefore, the highest possible score was 18. The task was stopped if four consecutive items were solved incorrectly. The score was used as the measure of non-verbal cognitive ability.

### Caregiver Questionnaires

Additionally, to better contextualise the environment in which the child was developing, caregivers completed questionnaires assessing home numeracy practices and family background.

Caregivers completed the Home-Math Environment Questionnaire (De Keyser et al., 2020). This measure inquired about math activities the child experiences at home. Out of 20 items in total, the first seven items measured the frequency of math-related activities at home (e.g., playing maths games or singing counting songs). The next ten items measured caregivers’ expectations regarding their child’s maths skills (e.g., hoping the child will master counting up to ten before starting school). The final three items measured caregivers’ attitudes towards maths (e.g. whether they consider maths skills important). The frequency of math-related activities at home sub-scale ranged on a 5-point Likert scale from 1 (never) to 5 (every day). Caregivers’ expectations were measured by a 4-point Likert scale from 1 (not at all important) to 4 (very important). Finally, caregivers’ attitudes towards mathematics were measured by a 5-point Likert scale from 1 (completely disagree) to 5 (completely agree). Scores for each subscale were computed as the average of the individual item responses.

We also asked caregivers to fill out a background questionnaire that included questions about the number of languages spoken in the household (Lefevre et al., 2002), the number of siblings (Wu et al., 2022), and the frequency of nursery attendance (Rajagopal et al., 2022). In addition, the questionnaire inquired about family income, the number of years spent in the UK, handedness, and relevant medical background (e.g., head injuries).

### fNIRS audio-visual matching task

A total of 36 audio-visual pairs were used in the audio-visual matching paradigm with three different tasks: easy numerical, difficult numerical, and non-numerical. Each task included 12 audio-visual pairs.

The easy numerical task contained pictures of one, two, and three apples. Auditory number words (“one”, “two”, “three”) were paired with the visual stimuli to create all possible audio-visual combinations. Matching trials (e.g., one apple paired with the number word “one”, two apples with “two”, and three apples with “three”) were each repeated twice to equate the number of matching and non-matching trials, resulting in six matching and six non-matching trials in total (see Figure 1A). Similarly, the difficult numerical task included pictures of five, six, and seven apples, which were paired with the corresponding auditory number words (“five”, “six”, “seven”) to create matching and non-matching trials. In the non-numerical task, images of an apple, banana, or orange were paired with auditory labels (“apple”, “banana”, “orange”) analogously. The voice recordings used were generally upbeat but not overtly child-directed, with a female native voice used for both English and Russian versions. Stimulus presentation was controlled by the OpenSesame software package (Mathôt et al., 2012). Tasks were presented in a fixed order: the easy numerical task, followed by the difficult numerical task, and then the non-numerical task. This order was installed to help children understand task structure and avoid early disengagement due to frustration with the difficult task. Appendix A contains the complete list of stimuli in the presentation order.

**Figure 1.**
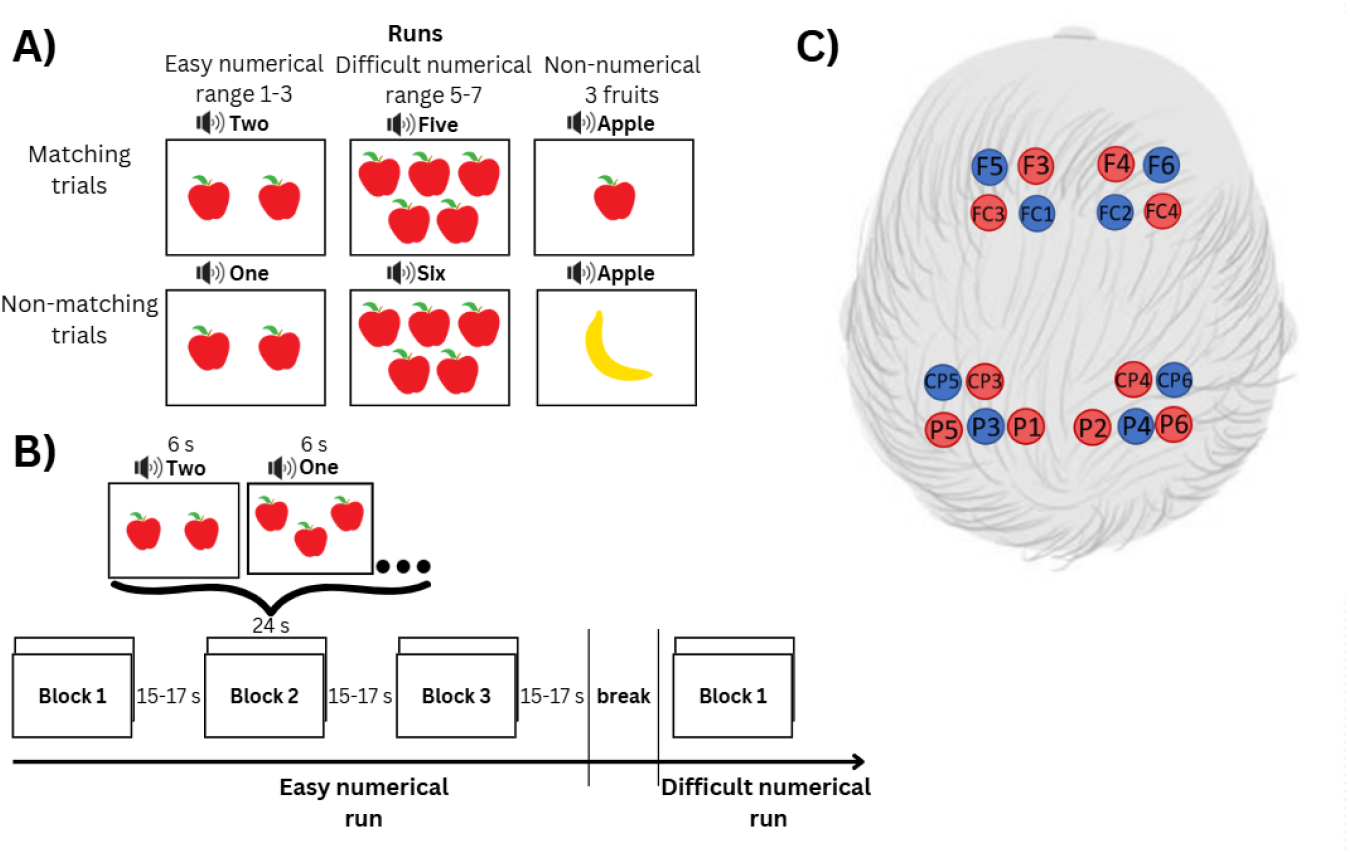
A) Task types. The easy numerical task contained pairs that ranged between numbers 1-3. The difficult numerical task had pairs that ranged between numbers 5-7. The non-numerical task had the same number of audio-visual pairs, and children were required to match the picture with the audio based on the semantic match, not the quantitative match. All tasks had six matching and non-matching pairs. B) The block design paradigm with an example of trials in one of the blocks. There were 3 blocks in one run. C) Positions of the fNIRS optodes based on the fNIRS Optodes’ Location Decider (fOLD) toolbox (Zimeo Morais et al., 2018). The red circles depict the sources, and the blue circles depict the detectors.

The task was conducted in three runs, each corresponding to one of the tasks (Figure 1B). Within each run, stimuli were presented in three blocks of four audio-visual trials in a pseudorandomised order. This block structure was chosen to maximise the signal-to-noise ratio of the fNIRS measurements, as longer blocks allow for more robust estimation of task-related haemodynamic responses compared with shorter blocks (Lloyd-Fox et al., 2010; Santosa et al., 2018). The duration of each audio stimulus was approximately 400 ms, varying slightly based on pronunciation time, and each trial allowed 6 s for the child to respond. The inter-trial interval was 500 ms. Each block lasted around 24 s, with jittered inter-block intervals of 15-17 s (mean of 16 s), during which a relaxing video of an aquarium was shown. The total duration of the experiment, including a self-paced break between tasks, was approximately 6 minutes. The task required a verbal “yes” or “no” response (i.e., determining whether a picture matched the audio) after each trial to ensure sufficient attention levels (Bugden et al., 2021; Kersey & Cantlon, 2017). For each task, performance was scored as the proportion of correct responses across trials.

### fNIRS recording

A portable continuous-wave fNIRS device (Brite, Artinis Medical Systems BV, The Netherlands) with 10 sources and 8 detectors with dual-wavelength (760 nm and 850 nm) near-infrared light was used to measure the absorption changes of oxygenated (HbO) and deoxygenated (HbR) haemoglobin (Maki et al., 1995). The data were recorded with a 25 Hz sampling rate, and the inter-optode distance was approximately 30 mm. The fNIRS cap, designed by Artinis Medical Systems BV, was arranged with 2×4 and 2×5 configuration of optodes, resulting in 18 measurement channels, with 9 channels on each hemisphere (Figure 1C). Emitters on the left and right parietal regions were placed on P3, CP5, P4, and CP6, respectively, with detectors positioned on CP3, P5, P1, P2, P6, and CP4. On the frontal regions, emitters were positioned on FC1, F5, FC2 and F6, respectively, with detectors placed on FC3, F3, F4, FC4. This arrangement adhered to the international 10-10 system (Chatrian et al., 1985).

### Data analysis

#### fNIRS preprocessing and analysis

The raw fNIRS data were pre-processed using the NIRS Brain AnalyzIR Toolbox (Santosa et al., 2018), an open-source, Matlab-based analysis package for managing, pre-processing, and performing both first- and second-level (i.e., single-subject and group-level) statistical analysis of fNIRS data. A custom pipeline was created for this pre-processing. The code is available via the following link: https://github.com/elizaveta-iva/Counting-apples.

After removing setup periods, we retained 5 s of pre- and 5 s of post-experiment baseline, and assessed data quality on a channel-by-channel basis using the QT-NIRS tool (Hernandez & Pollonini, 2020; Ivanova et al., 2026). QT-NIRS evaluates signal quality by examining factors like optical coupling between optodes and the scalp, motion artefacts, and saturated channels, based on the photoplethysmographic response related to the cardiac cycle. We followed the protocol of one of the few fNIRS studies in preschoolers that used QT-NIRS (Bulgarelli et al., 2023), for setting threshold values: a scalp coupling index (SCI) of ≥0.60 and peak spectral power (PSP) of ≥0.06. Both SCI and PSP were calculated for every 3-second non-overlapping window. Channels where both SCI and PSP fell below these thresholds for more than 60% of the windows were excluded from later analysis. On average, 3.52% of channels were excluded. Based on QT-NIRS assessments, all children had usable channels and were therefore included in further processing.

Next, the raw data were transformed into optical density data. We applied Wavelet motion artefact correction (Molavi & Dumont, 2012) with a threshold of 3 standard deviations and Temporal Derivative Distribution Repair (TDDR, Fishburn et al., 2019). The resulting signal of each fNIRS channel was converted into relative haemoglobin changes (HbO and HbR) using the modified Beer-Lambert law (Cope & Delpy, 1988), with a partial path length correction of 0.1 (differential path length factor = 6, partial volume factor = 60) for both wavelengths (Cope & Delpy, 1998). We then applied a fourth-order Butterworth filter with a band-pass range of 0.01-0.2 Hz and resampled the data to 2 Hz. A lower sampling rate was used for event-based analysis to reduce noise (Fishburn et al., 2014; Pinti et al., 2017).

Finally, we applied systemic artefact correction using an iterative autoregressive least-squares (AR-IRLS) implementation of the general linear model (GLM, Barker et al., 2013). Subject-level estimates were then entered into a mixed-effects model, with the task as a fixed effect and subject as a random effect, to obtain group-level statistics. Within each task, we identified significant beta values. Channels showing significant activation in the same direction (i.e., both increase or decrease) for both HbO and HbR were marked as noisy and not interpreted. Next, we ran separate t-tests corresponding to both HbO and HbR beta values to analyse contrasts of [difficult numerical - easy numerical task] and [easy numerical - non-numerical task]. Neurophysiologically, increased HbO and/or decreased HbR indicate increased brain activation (Fonseca et al., 2025; Sholkmann et al., 2014). Therefore, a channel was reported as significant only if it met one of two criteria: (i) a significant difference in either HbO or HbR, or (ii) both HbO and HbR were significant and changed in opposite directions. Multiple comparisons were corrected using the false discovery rate (FDR; q ≤ .05; Benjamini & Hochberg, 1995).

#### Exploratory connectivity analysis

To explore task-related interactions within the frontoparietal network, we computed functional connectivity for each task. Channels were grouped into four regions of interest (ROIs): bilateral frontal and parietal regions. For each task, we applied a robust autoregressive (AR) filtering model to the ROI time series to obtain temporally whitened signals. Connectivity was then estimated using robust AR-based correlations between these whitened signals, following procedures described by Fishburn et al. (2019). The resulting correlation coefficients were Fisher z-transformed (absolute values) to permit parametric statistical analyses. Finally, mixed-effects models were run for each ROI, with the task as a fixed effect and subject as a random effect. To aid interpretation, HbO-to-HbO and HbR-to-HbR correlations between the ROIs were analysed separately. Similarly to the contrast analysis, we reported FDR-corrected findings.

#### Behavioural analysis

To examine differences in performance accuracy across tasks, paired-samples t-tests were conducted to compare the difficult numerical vs. easy numerical tasks, as well as the easy numerical vs. non-numerical tasks. FDR correction was applied to account for multiple comparisons, and Cohen’s d was calculated to quantify the magnitude of effects.

#### Exploratory brain–behaviour analysis

To examine whether task-related neural responses were associated with behavioural performance, we conducted exploratory brain–behaviour correlations using accuracy measures. Analyses focused on channels that showed significant effects in the main contrast analyses. Pearson correlations were computed between channel-wise HbO and HbR responses and task accuracy, and results were evaluated with FDR correction for multiple comparisons.

## Results

### Behavioural results

#### Behavioural tasks and questionnaires

Children’s performance across behavioural measurements, as well as caregivers’ responses in questionnaires, is presented in Table 1.

**Table 1.**
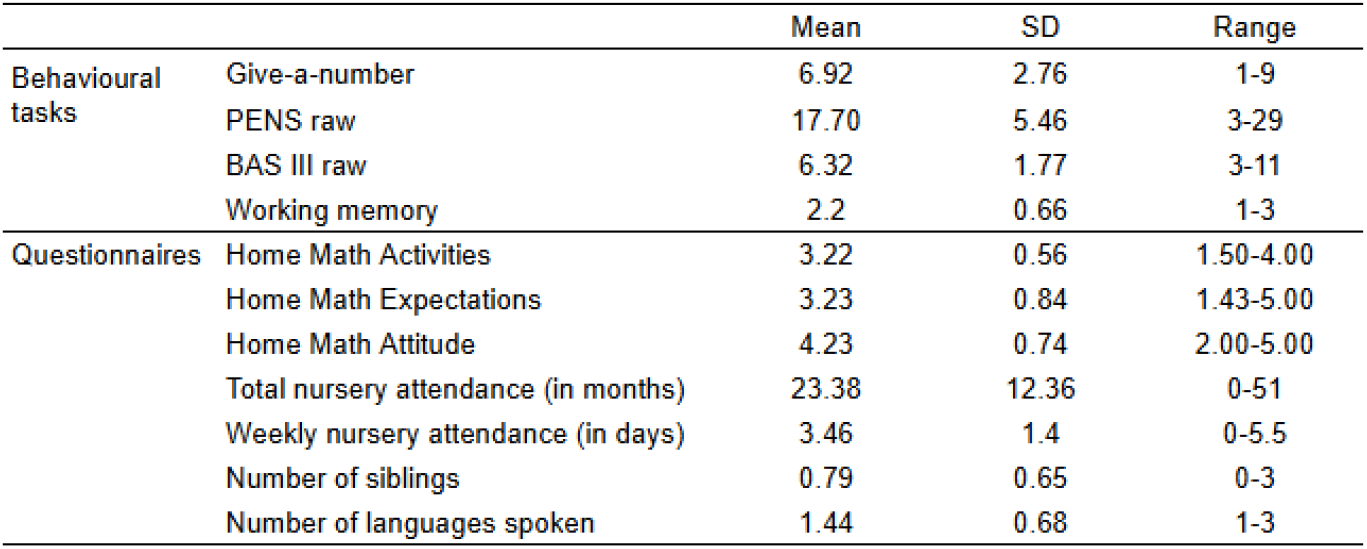
Demographics description.

#### Accuracy across the tasks

To examine differences in performance accuracy across tasks, paired-samples t-tests were conducted. Performance accuracy was significantly lower in the difficult numerical task (M = 0.58, SD = 0.23) compared with the easy numerical task (M = 0.93, SD = 0.16), t(59) = -11.65, p < .001, Cohen’s d = -1.50. Performance accuracy in the easy numerical task was significantly lower than in the non-numerical task (M = 0.98, SD = 0.08), although the effect was considerably smaller, t(59) = -2.65, p = .010, Cohen’s d = -0.34.

Given the relatively low performance in the difficult numerical task, a one-sample t-test was conducted to compare accuracy against chance level. Performance was significantly above chance, t(59) = 2.58, p = .012, Cohen’s d = 0.33.

### fNIRS results

#### Contrast analysis: Difficult numerical vs easy numerical

The t-test used to contrast the difficult numerical and easy numerical tasks showed increased hemodynamic response in the right middle frontal gyrus and right superior parietal lobule, as evidenced by an increase in HbO and a decrease in HbR, respectively. At the same time, the contrast elicited an opposite activation in the left inferior parietal lobule and left precuneus. These results suggest greater right fronto-parietal engagement during the difficult numerical task, but greater left parietal involvement during the easy numerical task (Table 2, Figure 2).

**Table 2.**
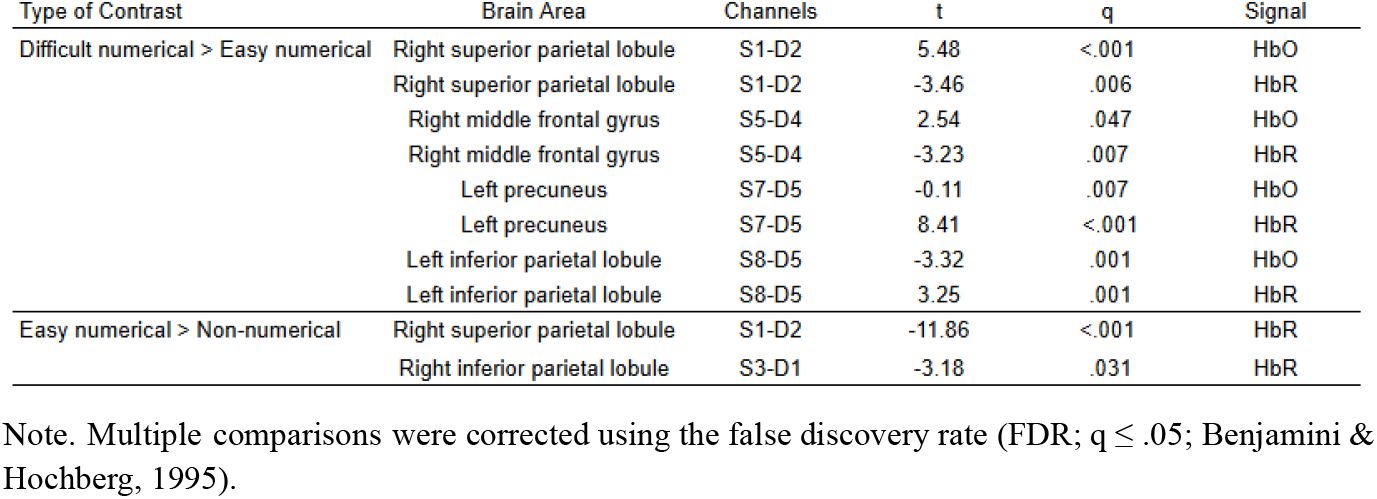
Summary of the significant channels in contrast analysis across the tasks.

| Type of Contrast | Brain Area | Channels | t | q | Signal |
| --- | --- | --- | --- | --- | --- |
| Difficult numerical > Easy numerical | Right superior parietal lobule | S1-D2 | 5.48 | <.001 | HbO |
|  | Right superior parietal lobule | S1-D2 | -3.46 | .006 | HbR |
|  | Right middle frontal gyrus | S5-D4 | 2.54 | .047 | HbO |
|  | Right middle frontal gyrus | S5-D4 | -3.23 | .007 | HbR |
|  | Left precuneus | S7-D5 | -0.11 | .007 | HbO |
|  | Left precuneus | S7-D5 | 8.41 | <.001 | HbR |
|  | Left inferior parietal lobule | S8-D5 | -3.32 | .001 | HbO |
|  | Left inferior parietal lobule | S8-D5 | 3.25 | .001 | HbR |
| Easy numerical > Non-numerical | Right superior parietal lobule | S1-D2 | -11.86 | <.001 | HbR |
|  | Right inferior parietal lobule | S3-D1 | -3.18 | .031 | HbR |

Note. Multiple comparisons were corrected using the false discovery rate (FDR; $q \leq .05$ ; Benjamini & Hochberg, 1995).

**Figure 2.**
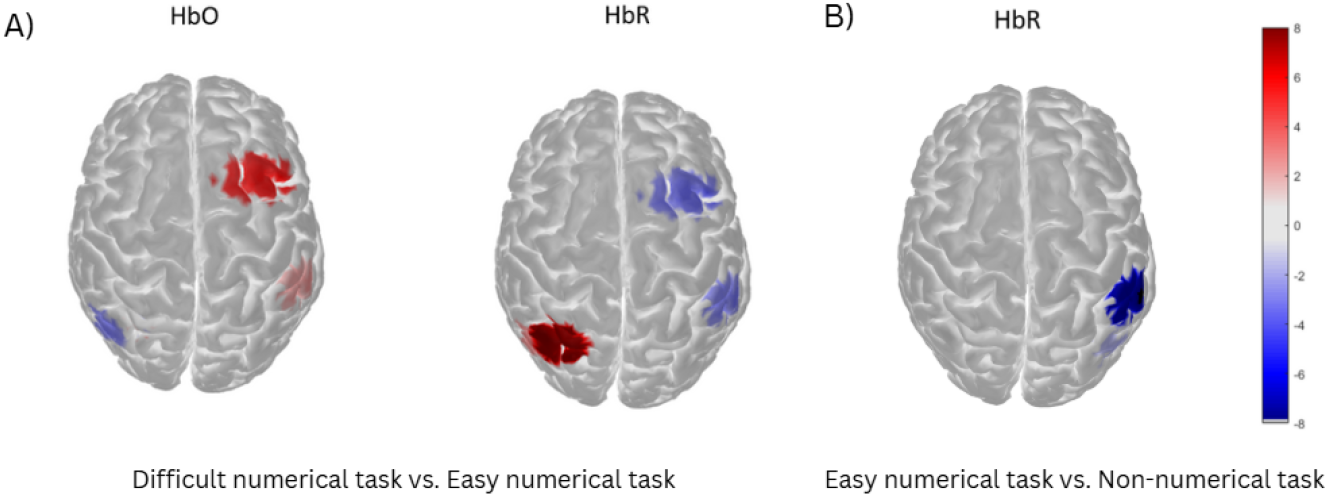
The heatmap of significant brain activation for the contrast of the difficult vs. easy numerical tasks (A) and easy vs. non-numerical tasks (B). Red and blue demonstrate increased and decreased activation in HbO, and the opposite pattern in HbR.

#### Contrast analysis: Easy numerical vs non-numerical

The contrast between the easy numerical and non-numerical tasks revealed greater activation in the right inferior and superior parietal lobules during the easy numerical task compared to the non-numerical task (Table 2, Figure 2).

### Exploratory analysis of the brain connectivity across tasks

The contrast between the difficult numerical task and the easy numerical task revealed increased connectivity between bilateral prefrontal regions in the easy numerical task (R=-.24, p=.02), albeit it did not survive FDR correction. There was no significant difference in connectivity between the easy numerical task and the non-numerical task. Additional separate analysis of each task showed that fronto-parietal connectivity emerged only under numerical tasks. Moreover, cross-hemispheric fronto-parietal links were observed exclusively in the easy numerical task (Table 3, Figure 3).

**Table 3.** Summary of the significant ROIs in connectivity analysis across the tasks.

| Condition | ROI A | ROI B | R | q | Signal |
| --- | --- | --- | --- | --- | --- |
| Non-numerical | Right Frontal | Left Frontal | .210 | .020 | HbO |
|  | Right Frontal | Left Frontal | .241 | .003 | HbR |
|  | Right Parietal | Left Parietal | .435 | <.001 | HbO |
|  | Right Parietal | Left Parietal | .389 | <.001 | HbR |
| Difficult numerical | Left Parietal | Left Frontal | .165 | .030 | HbO |
|  | Right Frontal | Left Frontal | .275 | <.001 | HbR |
|  | Right Parietal | Left Parietal | .539 | <.001 | HbO |
|  | Right Parietal | Left Parietal | .254 | .008 | HbR |
|  | Right Parietal | Right Frontal | .172 | .040 | HbR |
| Easy numerical | Left Parietal | Left Frontal | .155 | .047 | HbO |
|  | Right Frontal | Left Frontal | .379 | <.001 | HbO |
|  | Right Frontal | Left Frontal | .245 | .003 | HbR |
|  | Right Frontal | Left Parietal | .197 | .021 | HbO |
|  | Right Parietal | Left Parietal | .531 | <.001 | HbO |
|  | Right Parietal | Left Parietal | .407 | <.001 | HbR |
|  | Right Parietal | Right Frontal | .199 | .017 | HbO |

**Figure 3.**
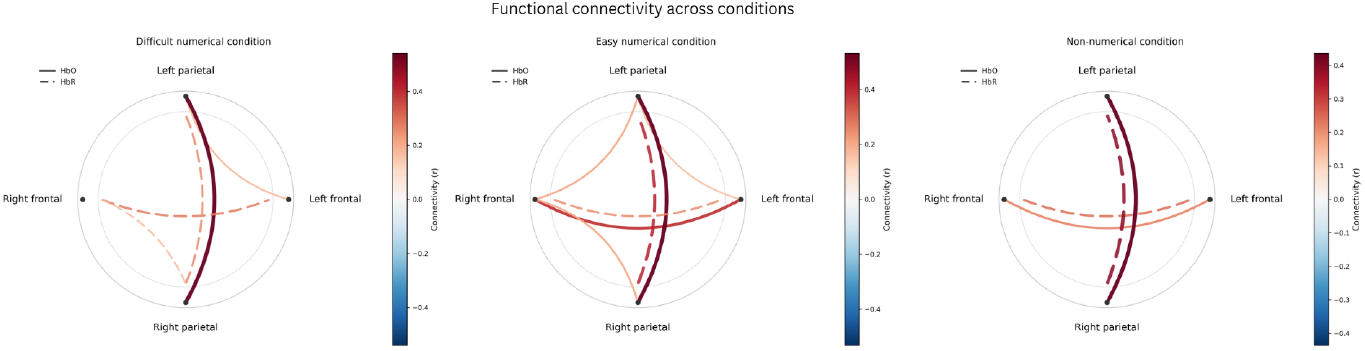
Connectivity heatmap across all tasks. Significant correlations (corrected for multiple testing, q<.05) are highlighted, while insignificant ones are left blank. For clarity, only separate HbO to HbO and HbR to HbR correlations are reported. In the non-numerical task, mixed-effects analysis revealed significant connectivity between the right and left frontal regions, as well as between the right and left parietal regions. In the difficult numerical task, we found significant connectivity between the right and left frontal and the right and left parietal regions. Additionally, there was a significant connectivity between left parietal and left frontal, and right parietal and right frontal regions. Lastly, in the easy numerical task, the mixed-effect analysis of connectivity revealed a series of significant connectivities between left and right frontal and left and right parietal regions. Similarly to the difficult numerical task, there was a significant fronto-parietal connectivity between left parietal and left frontal, left parietal and right frontal, and right parietal and right frontal.

### Exploratory analysis of the brain-behaviour correlations

#### Difficult numerical task vs. Easy numerical task

Overall, only one channel (S5-D4) in the right middle frontal gyrus, which was significant in the contrast analysis (see Table 2), was negatively correlated with the accuracy during the difficult numerical task (HbR: r=-.33, p=.038); however, this correlation did not remain significant after correcting for multiple comparisons.

#### Easy numerical task vs. Non-numerical task

No significant correlations were found between significant channels and accuracy during the easy numerical task after correcting for multiple comparisons.

## Discussion

In this study, we investigated the specificity of the prefrontal support for number processing in preschoolers. To do so, we asked children to match small (1-3) and large (5-7) numbers, as well as objects (fruits) to corresponding pictures, and recorded their brain responses using fNIRS. While our main focus was on prefrontal support, we also measured responses in the bilateral parietal regions, known to play a key role in number processing (Hyde, 2021; Skagenholt et al., 2025).

Our findings indicate that prefrontal engagement during early number processing is sensitive to task demands. Specifically, matching large numbers elicited greater activation in the right middle frontal gyrus compared to matching small numbers. Behaviourally, this condition was also more demanding, with an average accuracy of 58% for large numbers relative to 95% for small numbers. This interpretation is supported by previous research linking increased middle frontal gyrus activation to higher cognitive demands across a range of tasks in older children and adults (Ansari, 2005; Arsalidou, 2018; Battista et al., 2018; Bezuidenhout et a., 2026; Cantlon et al., 2009; Soltanlou et al., 2017). At the same time, the right lateralisation of middle frontal activation is notable, as this region has been associated with numerical processing in adults (Koyama et al., 2017). These findings suggest that while preschoolers rely on domain-general prefrontal support during particularly challenging numerical tasks, the lateralisation of this support may begin to align with regions that will later become specialised for numerical processing.

Our exploratory connectivity results support this interpretation. We observed both contra- and intralateral frontoparietal connectivity when children matched small numbers, but not when they matched large numbers or objects. The absence of frontoparietal connectivity during object matching is unsurprising, given the number-specific functionality of this network (Hyde, 2021). However, the presence of significant frontoparietal connectivity for small numbers, but not for large numbers, suggests that these networks are still undergoing functional development. Under these conditions, prefrontal engagement may support the coordination of domain-general executive processes when numerical knowledge is still developing (Best & Miller, 2010). This interpretation aligns with developmental models proposing that executive resources play a greater role in early symbolic number processing under high task demands, prior to the emergence of more efficient, specialised parietal representations (Ansari, 2012; Cantlon et al., 2009).

Even if there are signs of emerging number sensitivity within the prefrontal regions, the evidence for number-specific engagement should be taken with caution. Prefrontal activation did not differ between matching small numbers and non-numerical objects, suggesting that prefrontal responses were not uniquely driven by numerical content. A likely explanation lies in the developmental stage. Preschoolers in the present study are only beginning to acquire symbolic number knowledge (Sarnecka et al., 2015; Zhang et al., 2019), and the relevant frontoparietal networks may therefore still be in an early phase of functional organisation (Hyde, 2021). Consistent with this view, number-specific prefrontal responses have been reported more reliably in older children and adults, once symbolic numerical representations are more firmly established (Emerson & Cantlon, 2012, 2015).

Beyond prefrontal regions, we also observed distinct responses in parietal regions. When matching large versus small numbers, activation increased in the right superior parietal lobule. This pattern is consistent with prior work demonstrating both the distance effect, in which numerically close ratios (e.g., 6:7) elicit greater parietal activation than distant ratios, and the size effect, in which comparisons involving larger magnitudes (e.g., 6 vs. 7) recruit the right IPS more strongly than comparisons between smaller magnitudes (e.g., 2 vs. 3; Cantlon et al., 2006; Kersey & Cantlon, 2017; Pinhas et al., 2014). Although both effects are most often observed in non-symbolic numerical tasks (e.g., dot comparison), in our case, matching large numbers likely placed greater demands on maintaining and mapping symbolic representations to corresponding object sets. Given that the specialisation within the left parietal region was still ongoing, it is plausible that the more specialised right parietal region provided additional processing support in place. As such, our finding demonstrates a facilitatory role of the right parietal region for symbolic magnitude processing, in line with existing theories of parietal support for numerical cognition (Ansari, 2008; Kaufmann et al., 2011).

This interpretation is further supported by lower activation in the left inferior parietal lobule and left precuneus when matching large numbers relative to small numbers. Rather than indicating reduced involvement, this pattern might be due to a functional redistribution of processing demands, with the right parietal region increasingly supporting performance when symbolic processing becomes particularly challenging (Cantlon et al., 2010; Edwards et al., 2016; Hyde, 2021). Developmental evidence suggests that number-specific responses continue to strengthen across both hemispheres of the IPS throughout childhood, supporting bilateral processing (Emerson & Cantlon, 2015; Kaufmann et al., 2011).

Consistent with this view, the right inferior and superior parietal lobules showed greater activation when matching small numbers compared to objects. This pattern aligns with evidence implicating the right parietal cortex in magnitude processing across development, with robust numerical sensitivity observed in both children and adults (Cantlon et al., 2006; Kersey & Cantlon, 2017; Emerson & Cantlon, 2012). In contrast, we did not observe comparable specialisation in the left parietal region for this contrast. One possible explanation is that the left parietal cortex supports symbolic processing more broadly, rather than being selectively driven by numerical content (Ansari, 2008; Nieder, 2016), at least in preschoolers who are yet to fully develop symbolic numerical understanding. When symbolic processing demands are similar across numerical and non-numerical conditions, left parietal activation may therefore not differentiate strongly between them.

### Limitations and future directions

Our study has some limitations. First, we limited the experiment’s duration to accommodate young children’s low tolerance for wearing the fNIRS cap (Soltanlou et al., 2018). Consequently, we could not separately measure brain responses for matching and non-matching trials within each task. However, collapsing across tasks to compare matching versus non-matching trials may still provide insight into general error-detection processes (e.g., in prefrontal and anterior cingulate regions). Future research with designs optimised for trial counts could test this more directly and examine whether prefrontal responses differ between mismatched numbers and objects, providing a more detailed snapshot of frontoparietal activation. Similarly, we did not include a more difficult object-matching control, as the study primarily focused on numerical cognition. Comparing difficult numerical and non-numerical tasks in future research could help determine whether prefrontal specialisation is more attenuated when processing different types of challenging stimuli, though they may not overlap in number-specific regions (Chiou et al., 2023). Additionally, while fNIRS allows for flexible recording (Yucel et al., 2024), its spatial resolution is lower than that of fMRI, potentially missing activation changes in some prefrontal regions. Although an fMRI study could provide greater spatial precision, its feasibility with preschoolers is limited. Finally, we did not provide feedback on response accuracy during the task, and as such could not assess whether children perceived their responses as correct or incorrect. Such data could offer further insight into the mechanisms of prefrontal support, particularly by examining post-error enhancement of selective attention, which has been linked to increased prefrontal activation in adults (Weuthen et al., 2025).

## Conclusion

To our knowledge, this is the first study in preschoolers that tested both domain-general and number-specific mechanisms of prefrontal number processing. Our findings suggest that, unlike adults and older children, preschoolers appear to rely primarily on domain-general prefrontal support during symbolic number processing. This provides a neural framework for behavioural evidence demonstrating strong associations between domain-general abilities and early symbolic numerical knowledge, particularly in the preschool years (Emslander & Scherer, 2022), and stresses the importance of timely acquisition of these abilities during early learning. In addition, our study provides evidence for the supporting role of the right parietal region during more demanding symbolic numerical processing and suggests that the presence of frontoparietal connectivity may be associated with consolidated rather than acquired numerical knowledge.

## Acknowledgments

We thank the families who participated in our study. We also thank Dr Hadi Borj Khani for helping with the illustration of our neuroimaging findings and Robyn Copithorne Crainey for assisting with data collection.

## Data Availability Statement

The data that support the findings of this study are available on request from the corresponding author.

## Ethics Approval Statement

Written informed consent was collected from the caregivers. Additionally, each child was asked if they agreed to participate and could change their mind during the experiment. The procedures were accepted by the ethics committees at the University of Surrey (FHMS 22-23 073 EGA) and conformed to the requirements of the Declaration of Helsinki.

